# Lossless Pangenome Indexing Using Tag Arrays

**DOI:** 10.1101/2025.05.12.653561

**Authors:** Parsa Eskandar, Benedict Paten, Jouni Sirén

## Abstract

Pangenome graphs represent the genomic variation by encoding multiple haplotypes within a unified graph structure. However, efficient and lossless indexing of such structures remains challenging due to the scale and complexity of pangenomic data. We present a practical and scalable indexing framework based on tag arrays, which annotate positions in the Burrows–Wheeler transform (BWT) with graph coordinates. Our method extends the FM-index with a run-length compressed tag structure that enables efficient retrieval of all unique graph locations where a query pattern appears. We introduce a novel construction algorithm that combines unique ***`k`***-mers, graph-based extensions, and haplotype traversal to compute the tag array in a memory-efficient manner. To support large genomes, we process each chromosome independently and then merge the results into a unified index using properties of the multi-string BWT and r-index. Our evaluation on the HPRC graphs demonstrates that the tag array structure compresses effectively, scales well with added haplotypes, and preserves accurate mapping information across diverse regions of the genome. This indexing method enables lossless and haplotype-aware querying in complex pangenomes and offers a practical indexing layer to develop scalable aligners and downstream graph-based analysis tools. The index additionally supports efficient one-to-all coordinate translation, enabling any interval on a haplotype to be mapped to its corresponding intervals across all other haplotypes in the graph.

## 1 Introduction

Linear reference genomes act as essential blueprints in genomic research, offering a standard coordinate framework for comparing individual sequences [1, 2]. Aligning reads to this reference has enabled the discovery and interpretation of genetic variation across populations [3, 4]. Despite their utility, linear references come with significant drawbacks [5]. They represent only a single version of the genome and fail to capture the full extent of population-level variation [6]. This becomes particularly problematic in regions with complex structural variation or high diversity, where the reference may exclude certain sequences entirely or include only one of several possible configurations [7]. In such cases, reads from divergent haplotypes may align poorly, or not at all, resulting in ambiguous mappings and systematic bias in subsequent analyses called reference bias [8, 9].

To address these challenges, researchers have developed pangenome references, which aim to represent the full spectrum of human genetic diversity rather than a single canonical genome [10]. A popular usage of these references is to use them as graphs, where nodes represent sequences and edges capture possible continuations along different haplotypes [11]. By embedding multiple haplotypes and structural variants into a shared graph, pangenome representations can reduce reference bias and provide more accurate mapping for individuals whose genomes diverge from the traditional reference [12]. In recent years, graph-based representations have been adopted in several large-scale efforts to better characterize genomic variation across populations such as the Human Pangenome Reference Consortium (HPRC) and the African pangenome project [13–16].

Indexes that map sequences to their matching positions in the reference are essential tools for sequence alignment. In the common seed-and-extend approach, the index is first used to identify short exact matches, or seeds, between the query and reference. These seeds are then filtered and chained to provide a high-level structure of the potential alignment, which is finally refined using dynamic programming to obtain a full base-level alignment.

Several pangenome indexes have been proposed, but each comes with limitations. A space-efficient FM-index [17–19] can be constructed directly for the haplotype sequences. These indexes have been built efficiently for collections containing hundreds of human haplotypes [20–22], but they report the same seed separately for every haplotype in which it appears. Since aligners favor informative seeds with minimal redundancy, a method for merging equivalent seeds across haplotypes is required to make such indexes practically useful.

Alternatively, FM-indexes can be built directly for pangenome graphs [23–26]. As these indexes report seeds as graph positions, they effectively merge seeds at aligned haplotype positions. However, if the graph is not similar to a de Bruijn graph, index construction requires expensive graph transformations. These transformations can be lossy, meaning that the index will be missing some parts of the haplotypes. The construction process is also fragile, and it is not always possible to find suitable transformations for indexing the graph [23]. Despite the shortcomings, these graph indexes have been used in several read aligners [11, 27, 28].

Finally, minimizer indexes and other sparse *k*-mer indexes are the preferred approach in recent sequence aligners [29–31]. The index can be built quickly, and by making it report graph positions, we can avoid redundant seeds. However, the length of the seeds must be chosen in advance. This forces us to make a fixed trade-off between sensitivity and specificity that would not be required with FM-indexes.

The minimizer index used in the Giraffe read aligner [31] is an index of the haplotypes that reports the hits as graph positions. A recent theoretical work proposed using the same idea with FM-indexes, constructing a ”tag array” [32]. The FM-index is based on the Burrows–Wheeler transform (BWT), which is built by sorting the suffixes of the sequences in lexicographic order and listing the characters preceding each suffix in that order. If the sequences are similar, the characters preceding similar suffixes are likely the same. The BWT will then contain long runs of identical characters, making it highly compressible. By the same reasoning, if a pangenome graph is a reasonable alignment of similar sequences, the graph positions corresponding to similar suffixes are likely the same. A tag array that lists the graph positions for each suffix in lexicographic order should then be highly compressible. When we get a lexicographic interval corresponding to a pattern from an FM-index of the haplotypes, we can list the graph positions matching the pattern using document listing techniques [33, 34] over the tag array.

More recently, Olbrich and Ohlebusch [35] explored a related direction by computing TAG arrays over multiple sequence alignments. Their approach associates BWT positions with alignment columns and uses this structure to report distinct reference positions within FM-index intervals in an output-sensitive manner. Although their construction operates on aligned string collections rather than sequence graphs, it reflects a broader interest in practical TAG-based data structures and in using TAGs to map BWT intervals back to biologically meaningful coordinates.

Building on this theoretical foundation, we present the first practical implementation of tag array indexing for pangenome graphs. Our method uses a run-length compressed tag array to annotate FM-index intervals with graph positions, enabling efficient resolution of all occurrences of a query across haplotypes. Unlike previous approaches, it avoids redundant reporting and preserves full graph resolution without requiring lossy transformations. This indexing strategy can form the backbone of a scalable framework for haplotype-aware search and analysis over large pangenome graphs.

This is an extended version of a paper that was first presented in WABI 2025 [36]. In this journal version, we introduce several substantial additions and improvements. First, we implement an updated *k*-mer extension method that increases coverage and improves the robustness of the tag array construction. Second, we redesign both the r-index and tag array encodings to reduce disk usage and lower memory requirements during queries. Finally, we add a coordinate translation framework based on the tag arrays, enabling any haplotype interval to be mapped to all other haplotypes that follow the same path in the graph.

## 2 Preliminaries

### 2.1 Notation and background

Let Σ denote the DNA alphabet, and let *H* ∈ Σ*^n^* be a haplotype represented as a string of length *n*. For a collection of haplotypes H = *H*_0_*, H*_1_*, . . . , H_m−_*_1_, we define the concatenated text *T* = *H*_0_$_0_*H*_1_$_1_ *. . . H_m−_*_1_$*_m−_*_1_ where each $*_i_* is a unique endmarker not in Σ, and $_0_ *<* $_1_ *<* · · · *<* $*_m−_*_1_. This construction ensures that suffixes from different haplotypes are kept lexicographically separate.

For simplicity, let *N* = |*T* | and *T* [−1] = *T* [*N* −1]. The suffix array SA[0 *. . . N* −1] of *T* is an array such that *T* [SA[*i*] *. . .* ] is the *i*-th lexicographically smallest suffix of *T* . The Burrows–Wheeler Transform (BWT) of *T* is then defined as BWT[*i*] = *T* [SA[*i*] − 1]. All the end-markers in the BWT are considered to be the same symbol ”$”. The count array C is defined as C[*c*] = |{*j* | 0 ≤ *j < N* and *T* [*j*] *< c*}| for each symbol *c* ∈ Σ∪{$} which is the total number of characters in *T* that are strictly smaller than *c*.

The rank function rank_BWT_(*c, i*) gives the number of occurrences of character *c* in the prefix BWT[0 *. . . i* − 1], that is rank_BWT_(*c, i*) = |{*j* | 0 ≤ *j < i* : BWT[*j*] = *c*}|.

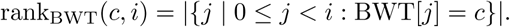

The multi-string BWT (MSBWT) is the Burrows–Wheeler Transform of a collection of strings, formed by concatenating the individual haplotype sequences with distinct end-markers as described above. By assigning a unique end-marker to each haplotype, the MSBWT ensures that no suffix of one haplotype overlaps lexicographically with suffixes of others. This construction guarantees that the suffix array SA of the concatenated text *T* respects haplotype boundaries and that the resulting BWT maintains a consistent ordering across the entire collection.

Let Tag[0 *. . . N* − 1] be an array such that Tag[*i*] stores the label associated with the suffix *T* [SA[*i*] *. . .* ]. In our setting, this label corresponds to the graph position from which the suffix originates. Each graph position is defined by a triplet (v, o, b), where v is the node identifier, o is the offset within the node, and b ∈ {0, 1} indicates whether the position lies on the reverse strand. The tag array thus provides a positional annotation over the BWT, enabling the recovery of graph coordinates from lexicographic intervals. Figure 1 shows a toy example of the graph and an illustration of the tags based on their positions on the graph.

**Fig. 1:**
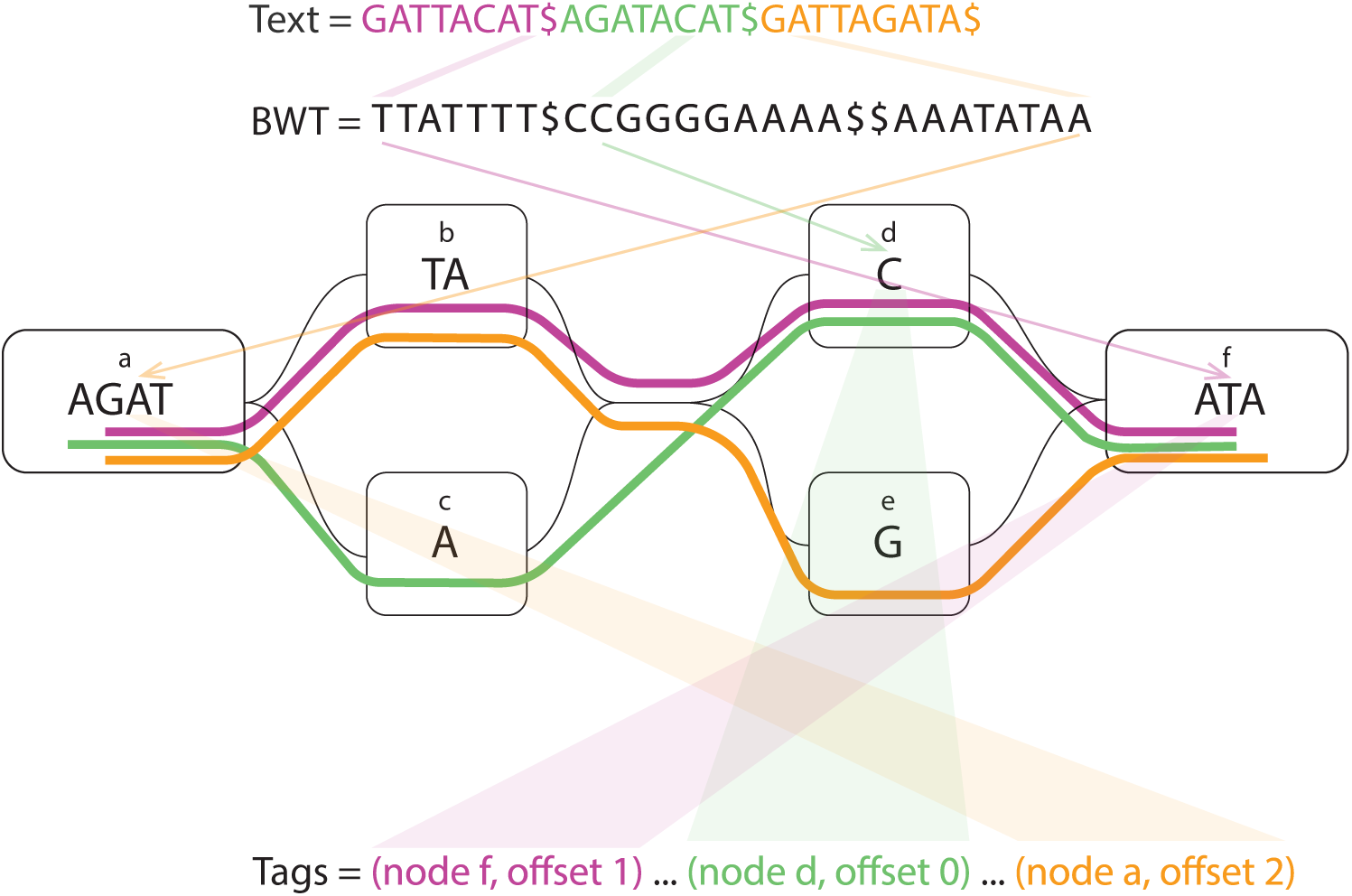
Toy example of the graph and tags. The haplotypes on the graph are shown with purple, green, and orange stripes.

### 2.2 LF-mapping and the FM-index

The FM-index [17] is a compressed full-text index built on the BWT. It supports efficient pattern matching by performing backward search. Instead of scanning the text directly, it iteratively refines the range of suffixes that match a given pattern, one character at a time, from right to left. This operation is using the Last-to-First (LF) mapping, which is central to the FM-index.

LF-mapping enables traversal of the BWT in a manner that corresponds to moving backward in the original text. It maps a position *i* in the BWT to the position in the suffix array where the suffix that precedes SA[*i*] begins. There are two common ways to express the LF-mapping:

1. LF-mapping with an arbitrary character *c*:

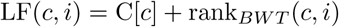

which computes the LF-mapping for character *c* at position *i* in the BWT.

2. LF-mapping using the character at position *i* in the BWT:

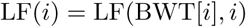

This is the standard LF-mapping, where we use the actual character occurring at position *i* in the BWT.

Using the LF function, the FM-index can search for a query pattern *P* = *p*_0_*p*_1_ · · · *p_m−_*_1_ by processing the characters from right to left. This process iteratively narrows a range [*A . . . B*] in the suffix array such that all suffixes in this interval begin with *P* . This is known as the find operation and is used to identify the lexicographic interval that matches the pattern.

To determine the exact positions where the pattern occurs in the original text, the locate operation is used, which recovers the corresponding suffix array values within the interval. This typically relies on a sampled suffix array, where positions are stored at regular intervals that control the sample rate [17, 37]. Locating a match that does not fall on a sampled position requires following a sequence of LF steps backward until a sample is reached. This creates an inherent time-space trade-off: denser sampling enables faster location but increases memory and disk usage. In highly repetitive texts such as pangenomes, locating can become inefficient, as the trade-off does not improve with the compressibility of the texts.

### 2.3 FMD-index

The FMD-index [38] extends the FM-index to support bidirectional pattern search by indexing both a text and its reverse complement in a unified data structure. While bidirectional pattern extension was previously introduced using paired BWTs [39], the FMD-index simplifies this approach by combining the forward and reverse indexes and incorporating reverse complements directly. This design enables efficient matching of a query sequence to either strand of the reference without requiring orientation-specific preprocessing.

The FMD-index is particularly useful in read mapping, where the orientation of the read relative to the reference is not known in advance. Its bidirectional search capabilities also allow it to support algorithms for finding maximal exact matches (MEMs), which serve as informative seeds for downstream alignment. The forward-backward search algorithm introduced in [38] enables the enumeration of all MEMs that contain a given position in the query, and this approach has since been adopted in popular tools such as BWA-MEM [40]. More recently, the algorithm was made faster by avoiding short MEMs that are unlikely to be relevant [41].

### 2.4 r-index

When the text is highly repetitive, the FM-index can be compressed well by run-length encoding the BWT [18, 42]. However, the usual approach for locating the occurrences of the pattern does not work well with highly repetitive texts. We have to make an unattractive trade-off between slow queries and using much more space than the rest of the index.

The r-index [19] solved the problem of locating the occurrences. It stores suffix array samples at run boundaries, making the space usage scale with the number of runs in the BWT. With some additional structures, it can derive SA[*i* + 1] from SA[*i*] (or the other way around). By starting from a run boundary or from a toehold found during pattern matching, we can report a large number of occurrences efficiently.

## 3 Methods

Our method consists of two main components: a construction phase, in which the tag arrays are built over the pangenome graph using a combination of unique *k*-mers, extension, and traversal; and a query phase, in which the resulting tag arrays are used to efficiently extract graph-level information for given input patterns. In addition, we introduce a coordinate-translation mechanism that translate coordinates from a chosen haplotype to their equivalent positions across all other haplotypes.

### 3.1 Construction

We build the tag array index using a multi-stage construction algorithm. Naive approaches, such as traversing every haplotype or assigning tags in text order and permuting with the suffix array, require storing large arrays in memory. This is not feasible for large pangenomes. For example, the final stage of our algorithm, in which we traverse all the paths in order to cover all BWT positions, could build the tag array on its own. But because traversing the haplotypes in text order corresponds to filling the tag array in an arbitrary order, there would be a large number of short runs and short gaps halfway through the construction.

Instead, we use a more memory-efficient strategy. Our method combines a run-length encoded B+ tree — a self-balancing tree — with a layered construction pipeline. We first use unique *k*-mers as anchors to annotate the BWT. Then we extend these *k*-mers to increase coverage. Finally, we traverse each haplotype to fill in the remaining gaps. To reduce memory usage and improve scalability, we perform this process separately for each chromosome as shown in Figure 2. At the end, we merge the tag arrays across chromosomes into a single global index by using the structure of the multi-string BWT and the BWT index to sequence number mapping provided by the r-index.

**Fig. 2:**
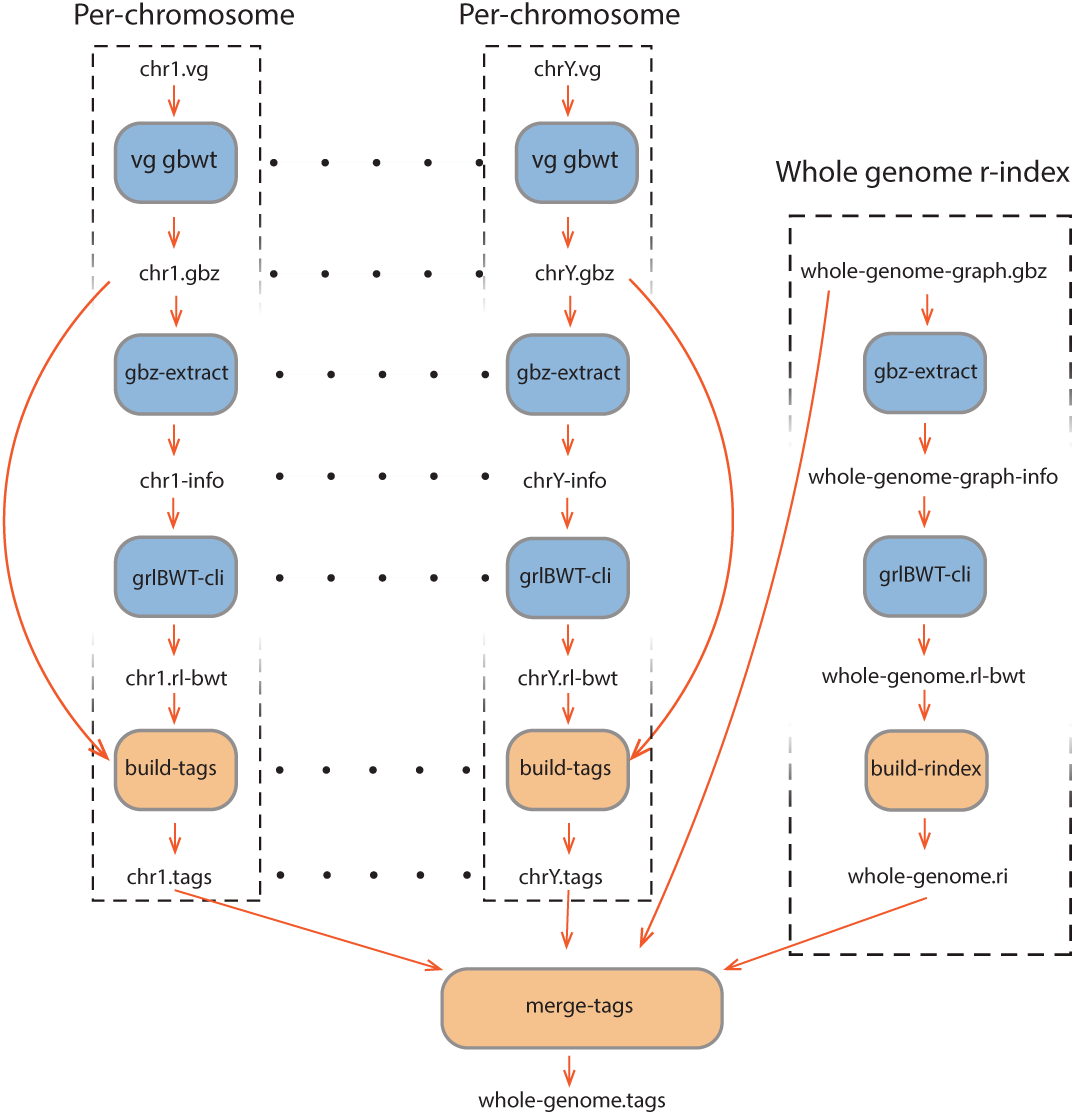
Tag array index construction workflow. Blue boxes are the external tools used in the pipeline, and orange boxes are the internal functionalities.

#### 3.1.1 Run-length B+ tree for efficient tag array construction

To efficiently manage the runs of tags during tag array construction, we designed and implemented a Run-Length B+ Tree (RLB+). This data structure extends the traditional B+ tree by incorporating mechanisms for handling run-length encoding of tags, enabling efficient storage, insertion, and merging of runs. In our context, a run of tags is defined as a contiguous segment of the tag array where all positions correspond to the same value (tag), which originates from the same position in the pangenome graph. Each run is stored as a triplet consisting of a key (tag), a start position, and a run length. This structure allows us to gradually compute tags and merge them efficiently. Unlike standard tree-based indexes that store individual keys, the RLB+ captures longer homogeneous segments compactly, leading to substantial space and time efficiency during tag propagation and merging.

Each leaf node in the RLB+ stores up to a fixed number of tag runs, determined by the degree of the tree. Each run consists of a BWT start position and a corresponding graph position. Instead of explicitly storing the run lengths, they are implicitly derived from the difference between the start positions of consecutive runs within the leaf or across adjacent leaves. In cases where there is no adjacent run following the current one, we insert a sentinel key at the final BWT position, using a reserved graph position to mark the end. This compact representation reduces memory overhead while maintaining efficient search and update performance.

Insertion into the RLB+ follows the standard B+ tree insertion procedure, but with additional logic to handle run merging. When a new tag run is added, the tree determines whether it can be merged with an adjacent run based on the graph position:

1. No Merge Condition: If the new run does not share the same graph position as the adjacent runs, it is inserted as a separate entry.
2. Forward Merge: If the new run extends an existing run (i.e., it has the same graph position as the previous run), it is merged with the preceding run.
3. Backward Merge: If the new run follows an existing run with the same graph position, it is merged with the succeeding run.
4. Bidirectional Merge: If the new run bridges two adjacent runs with the same graph position, all three segments are merged into a single, larger run.

A key distinction between the RLB+ and a standard B+ tree is its handling of node underflow. In a conventional B+ tree, the insertion operation ensures that underflow never occurs at any level of the tree. However, in the RLB+, the merging of runs introduces a scenario where underflows may arise. Specifically, in a bidirectional merge, the number of keys in a leaf node is reduced, potentially causing the node to fall below the minimum key threshold. To handle this, the RLB+ implements a rebalancing mechanism that ensures the structural integrity of the tree while preserving efficient search and update operations. When an underflow is detected in a leaf node:

- The tree first attempts to borrow a key from a neighboring sibling, maintaining balance while avoiding additional structural changes.
- If borrowing is not possible, a leaf merge operation is triggered, combining the underflowing node with its adjacent sibling and adjusting the parent accordingly.
- If the underflow propagates upward due to excessive merging, the internal nodes are recursively adjusted, following the standard B+ tree balancing rules.

By integrating run merging with dynamic rebalancing, the RLB+ maintains its logarithmic time complexity for insertions and queries while efficiently managing run-length encoding in the tag array. This makes it particularly well-suited for large-scale pangenome indexing tasks, where compact and dynamic storage of tag information is essential.

#### 3.1.2 Extracting tags from unique k-mers

The first step in constructing the tag array involves using the unique *k*-mers of the pangenome graph as shown in 3(A). A unique *k*-mer is a substring of length *k* with exactly one starting position in the pangenome graph. By identifying these unique *k*-mers, we establish anchors that allow us to map intervals of the BWT to graph positions.

Given a set of unique *k*-mers and their corresponding graph positions, we use the r-index to find these *k*-mers in the BWT. The r-index efficiently supports LF-mapping, allowing us to compute the suffix array interval [*A . . . B*] for each unique *k*-mer. Since these *k*-mers are unique in the graph, all occurrences in the BWT must correspond to the same graph position. This results in a run-length encoding of the tag array, where:

- The starting position of the run is defined as *A*, which is the starting position of the BWT interval [*A . . . B*].
- The graph position of the unique *k*-mer determines the tag assigned to the run.
- The run length is computed as *B* + 1 − *A*, covering all occurrences of the *k*-mer in the BWT.

Each such run is stored in the RLB+, ensuring efficient insertion, merging, and retrieval. This representation captures the mapping between suffix array positions and graph positions.

#### 3.1.3 Extending unique k-mers

After identifying the unique *k*-mers in the pangenome graph and mapping their corresponding BWT intervals, the next step is to extend these *k*-mers to maximize the coverage of the tag array. If *k*-mer *U* is graph unique and the graph position has only one predecessor with character *c*, then the (*k* + 1)-mer *cU* is also graph unique. So, we can backward-extend those unique *k*-mers with the additional bases as shown in with an example in 3(B).

For a unique *k*-mer *U* positioned at a specific node in the graph, extension is performed as follows:

1. Graph-based Backward Extension: If the preceding base *c* in the graph does not introduce new variants, either belongs to the same node or has only one predecessor node, the *k*-mer is extended one base at a time to *cU* , until a variation or a graph boundary is encountered.
2. In addition to extending only through unique predecessors, we also generalize the extension rule to handle predecessor sets that contain variation. When multiple predecessor nodes exist, we group them by the character of their last base. If a character appears in only one predecessor node, that branch is effectively unique for that character, and we extend the *k*-mer through it. For example, if the five predecessors nodes end in A, A, A, T, and C, we extend through the T and C nodes, as each base is contributed by exactly one predecessor and therefore represents an unique continuation.
3. BWT Interval Update via LF-Mapping: Since the BWT interval for *U* is already known and stored in the RLB+, the interval for the extended (*k* + 1)-mer *cU* can be efficiently computed using LF function by backward extending *U* with *c*.
4. Tag Array Expansion: We compute the extended (*k* + 1)-mer graph position using the original unique *k*-mer, and its new BWT interval is added to the RLB+.

By iteratively extending unique *k*-mers, a larger fraction of the tag array is populated, ensuring that we have fewer positions to fill in the final stage.

#### 3.1.4 Filling the gaps using haplotype traversal

Despite the substantial coverage obtained from unique *k*-mers and their extensions, a significant fraction of the tag array remains uncovered, particularly in regions that do not contain uniquely identifiable *k*-mers. To resolve this, we introduce a final step in the construction of the tag array that ensures complete coverage by leveraging the known haplotype information embedded in the pangenome graph.

Because we operate on the MSBWT, each haplotype corresponds to a unique string in the BWT. For each haplotype, we identify its endpoint in the BWT using the properties of the suffix array and the MSBWT. Once the BWT endpoint of a haplotype is known, we perform a backward traversal along the sequence of that haplotype, effectively walking in reverse from the end of the sequence to its beginning.

At each step in the backward traversal:

1. We determine the BWT position of the current character using the backtracking function.
2. The graph position associated with the current haplotype is known from its sequence in the graph.
3. We search the RLB+ to check whether this BWT position has already been assigned a tag. If it has, we continue traversal without modification.
4. If no tag exists for that position, we insert a new run in the RLB+ with the current BWT position and the graph position of the haplotype at that point.

This traversal is repeated independently for each haplotype, ensuring that every BWT position is assigned a tag. By filling in all previously unassigned regions of the tag array, this step guarantees that the entire BWT is labeled.

#### 3.1.5 Per-chromosome tag array construction

Applying tag array construction across an entire pangenome is computationally demanding due to the vast size and complexity of multigenome references. Constructing the tag array over the complete pangenome graph at once requires extensive memory resources and may not scale efficiently. To address this, we adopt a modular strategy by computing the tag arrays independently per chromosome.

In this approach, the tag array construction algorithm is applied separately to the subgraph corresponding to each chromosome. Each graph at chromosome level is processed independently, allowing the use of more manageable memory footprints while enabling parallelism across chromosomes. The output of each run is a tag array specific to that chromosome, representing the association between the BWT intervals and the graph positions for the sequences contained in that subgraph.

#### 3.1.6 Merging per-chromosome tag arrays into a whole-genome index

After computing the tag array indexing separately for each chromosome, we obtain localized tag arrays that map BWT positions within each chromosome to their respective graph positions. However, for downstream applications—such as whole-genome querying and alignment—it is necessary to combine these chromosome-specific tag arrays into a single global tag array indexed over the full pangenome graph.

Multi-string BWTs can be merged by constructing an interleaving array that maps how suffixes from individual texts (or chromosomes) should be ordered in the global BWT [43, 44]. In our approach, we use the same properties for interleaving the chromosome-specific tag arrays into a whole-genome tag array. We generate the interleaving array on the fly by iterating over the suffix array using a whole-genome r-index with multiple threads. The same r-index will also be used for queries.

The key challenge in this step is determining, for each BWT position in the whole MSBWT, which chromosome it belongs to, so we can read from the tag arrays of that specific chromosome. We use the structural properties of the MSBWT, in which the text is formed by concatenating all sequences (haplotypes) with unique end-markers. As a result, for every position in the BWT, it is possible to determine the sequence number—i.e., the index of the original haplotype to which that suffix belongs.

To bridge from sequence numbers to chromosome identifiers, we use the graph topology encoded in the GBZ format of the pangenome [45]. Specifically, we compute the weakly connected components of the graph, each of which corresponds to a distinct chromosome. Using the graph, we can easily determine the related chromosome for each of the sequence numbers. Then by inspecting just one graph position in each of the per-chromosome tag arrays we can determine the mapping between the components of the graph and the chromosomes.

With this mapping in place, we can interleave the chromosome-specific tag arrays into a global tag array, where each BWT position is assigned the tag value from its corresponding per-chromosome tag array. This construction preserves the correct positional and annotation semantics of each tag while enabling unified, whole-genome indexing and query capability.

### 3.2 Querying the tag array index

The other component of our method is an efficient query interface that returns the set of unique graph positions (tags) corresponding to any substring pattern in the pangenome. This enables applications such as haplotype-aware read mapping.

Given a query string P, we use the r-index to find its lexicographic interval [*A . . . B*] in the whole genome BWT of the haplotypes. This interval corresponds to all suffixes of the reference that begin with P. Our goal is to identify all distinct tags that annotate the BWT positions within this interval.

#### 3.2.1 Data structure

Instead of using the RLB+ for queries, we opt for a simple immutable structure that is both smaller and faster. The tag array is stored in a run-length encoded form to reduce space usage. Specifically, we store a sparse bitvector using Elias–Fano encoding [46] which marks the beginning of each tag run, and a vector of tags corresponding to each run.

For a sequence of tag runs [(*T*_1_*, L*_1_), (*T*_2_*, L*_2_), (*T*_3_*, L*_3_)*, . . .* ], where each *T_i_* is a tag and *L_i_* is the length of the run, the bitvector contains the positions [0*, L*1*, L*1+*L*2*, L*1+ *L*2 + *L*3*, . . .* ]. Each value in this vector marks the starting BWT position of a tag run, and the corresponding tag *T_i_*is associated with the interval between consecutive positions. This allows us to identify the tag for any given BWT position using rank-based queries on the bitvector.

#### 3.2.2 Query algorithm

To query which tags are present in the BWT interval [*A . . . B*], we perform two rank queries on the bitvector. One at position A returns the start run, and one at position B returns the end run.

The tag values for the runs overlapping [*A . . . B*] are then collected from the tag vector. We return the set of distinct tags by sorting the tags and removing the duplicates. These tags are associated with all runs in the range from the start run to the end run. This operation is efficient and avoids the need to scan or decompress the full tag array.

### 3.3 Coordinate translation using sampled tag arrays

In addition to pattern-to-graph queries, we support one-to-all coordinate translation between haplotypes using a compact sampled tag array built on top of the full tag array index. The goal of the translation query is the following: given a haplotype and an interval on that haplotype, report all other haplotypes and offsets that pass through exactly the same graph positions.

#### 3.3.1 Sampled tag arrays

We implement the sampled tag array by storing only those tag runs whose graph offset is *o* = 0, that is, the tag values marking the beginnings of nodes in the graph. Each such sampled tag is stored in the order of its corresponding BWT position in a wt gmr data structure [47]. Conceptually, the sampled tag array consists of alternating sampled tag runs (at node boundaries) and gap runs (covering all interior offsets), which makes it straightforward to determine whether a BWT position corresponds to a sampled tag or a gap. The wt gmr structure lets us retrieve the tag value of any sampled run and enumerate all sampled BWT positions that carry a given tag value. Because sampled tags exist only at node boundaries while all interior offsets are treated as gaps, this representation is far smaller than the full tag array yet remains sufficient for propagating coordinates along shared graph paths.

#### 3.3.2 Coordinate translation query

A coordinate translation query takes as input a sequence identifier sid and an interval [*p, q*) on that sequence, and returns all pairs (sid*^′^, p^′^*) such that the haplotype sid*^′^* passes through the same graph positions as sid over that interval. The query is implemented in two phases: (1) finding the set of tags traversed by the input interval, and (2) finding all haplotypes and offsets that traverse those tags.

In the first phase, we map the input interval [*p, q*) on sequence sid to BWT positions using the r-index. Starting from the end of the interval, we locate the BWT position corresponding to (sid*, q* − 1) and then iterate backwards towards (sid*, p*) with LF-mapping. Whenever we encounter a sampled position *i*, we retrieve its tag from the sampled tag array and add it to the set of tags traversed by the query interval. This phase yields the set of node-start tags that the input haplotype interval passes through. In the second phase, we translate these tags to coordinates on all other haplotypes.

For each tag *T* collected in the first phase, we use the wt gmr structure to enumerate all sampled positions whose tag is equal to *T* . Each such occurrence corresponds to a tag array run and determines a BWT interval where the underlying graph position is fixed to *T* . We then use the r-index to locate all pairs (sid*^′^, p^′^*) that fall into that BWT interval, obtaining all haplotypes and offsets that pass through the same graph position. Finally, we aggregate the results over all tags and report, for each tag, the offset of the tag within the query interval on sid together with the corresponding (sid*^′^, p^′^*) pairs on other haplotypes.

This two-phase procedure produces a one-to-all mapping from a haplotype interval to all other haplotypes that share the same path through the graph over that interval, while leveraging the compressed sampled tag array and r-index to keep both time and space usage practical at pangenome scale.

## 4 Results

### 4.1 Experimental setting

In all experiments, we use the bidirectional index of our method, built on both the forward and reverse strands of the pangenome graphs by constructing the FMD-index. This bidirectional representation enables applications such as MEM finding and other algorithms that rely on extending queries in both directions. However, it also doubles the total number of bases and increases the number of runs in both the BWT and the tag array compared to a unidirectional index. We further discuss the implications of this bidirectional index in Section 4.6.

We conducted our experiments on the Phoenix compute cluster at the University of California, Santa Cruz. Each node had dual AMD EPYC 7662 64-core processors, with 64 physical cores per processor and two hardware threads per core, yielding a total of 256 logical CPUs. The system had 2.0 TiB of RAM. Although the node was part of a shared cluster, all runs were performed within exclusive SLURM job allocations, ensuring that no other user processes interfered with our computations during runtime.

We used two human pangenome graphs from the Human Pangenome Reference Consortium (HPRC) [13]. A detailed summary of the properties of the graphs is provided in Table 1, including the number of haplotypes, total sequence length, and BWT and tag array statistics. See Appendix A for further details.

**Table 1:**
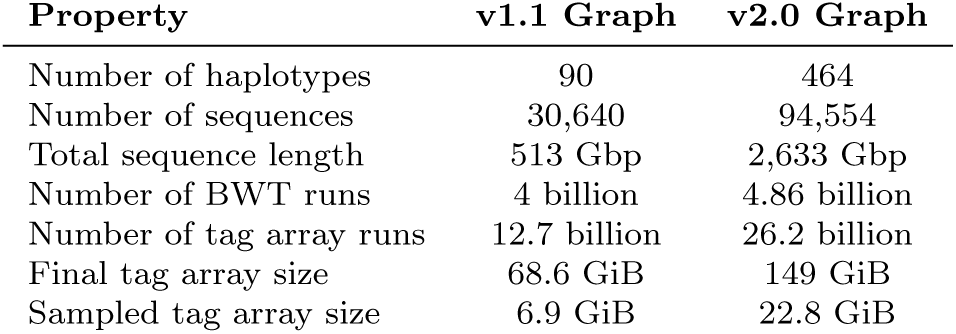
Summary statistics of the two HPRC pangenome datasets. These statistics correspond to the bidirectional version of the index, where both forward and reverse strands of all pangenome haplotypes are included.

| Property | v1.1 Graph | v2.0 Graph |
| --- | --- | --- |
| Number of haplotypes | 90 | 464 |
| Number of sequences | 30,640 | 94,554 |
| Total sequence length | 513 Gbp | 2,633 Gbp |
| Number of BWT runs | 4 billion | 4.86 billion |
| Number of tag array runs | 12.7 billion | 26.2 billion |
| Final tag array size | 68.6 GiB | 149 GiB |
| Sampled tag array size | 6.9 GiB | 22.8 GiB |

### 4.2 Performance of index construction

For building the tag array index, we used the grlBWT tool [21] to build the run-length encoded BWT. We built our r-index implementation on the basis of the compact output from the grlBWT. Figure 4 compares the time and memory usage of the tag array construction. The time for building the BWT using grlBWT is not included in this figure. The maximum time for building the BWT for a chromosome of the v1.1 graph was 35 minutes, and for the v2 graph was 75 minutes, all using 16 threads. The maximum total elapsed time for per-chromosome tag arrays building, including the BWT construction, was 6.7 hours for the v1.1 graph, and 26.5 hours for the v2 graph.

**Fig. 3:**
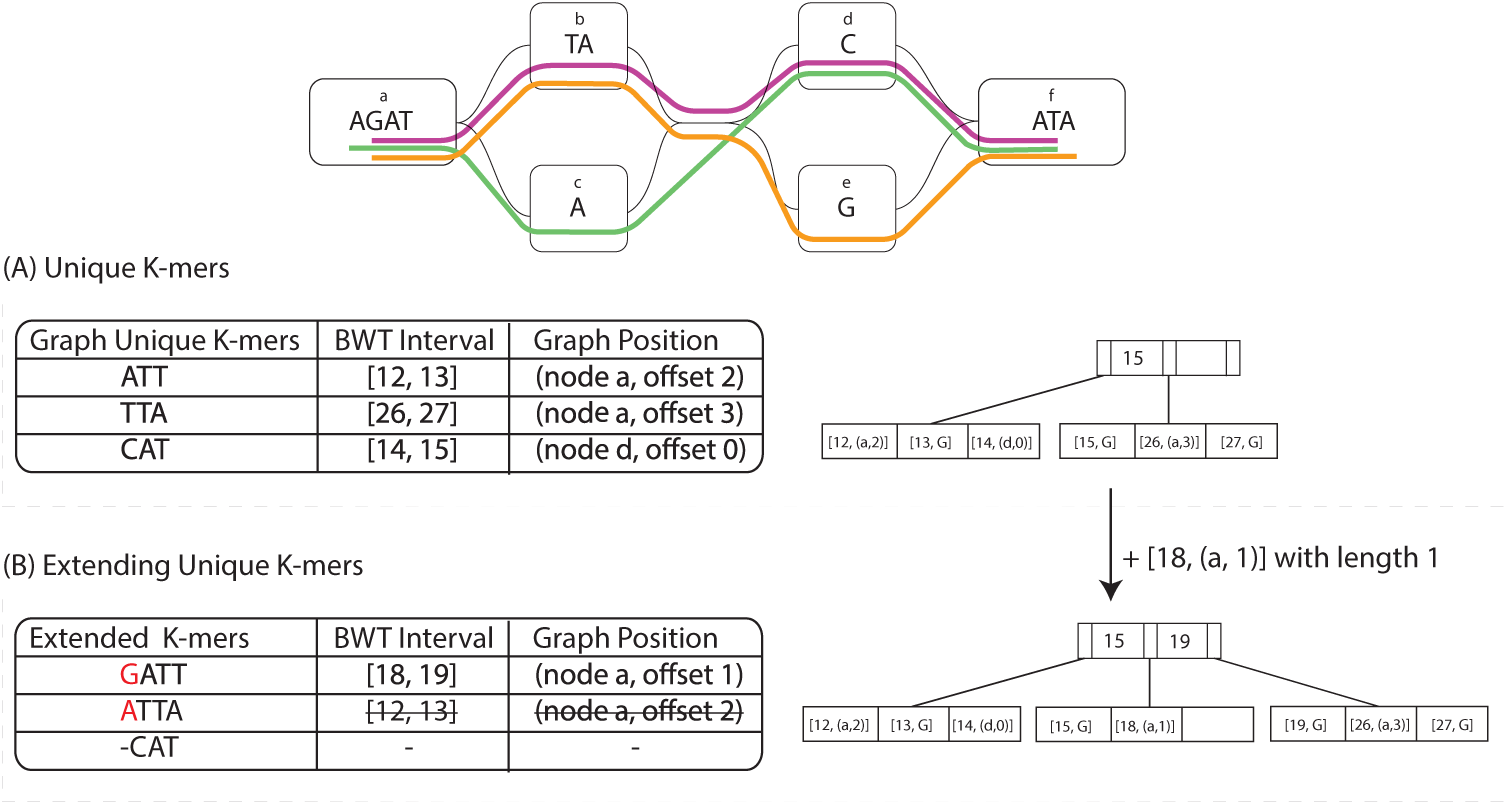
An example of the building tag array algorithm. (A) shows the unique 3-mers of the toy graph and their BWT intervals and graph positions on the left, and the structure of the RLB+ with the data on the right side. (B) show one step of extending the unique 3-mers and the RLB+ structure after that extension.

**Fig. 4:**
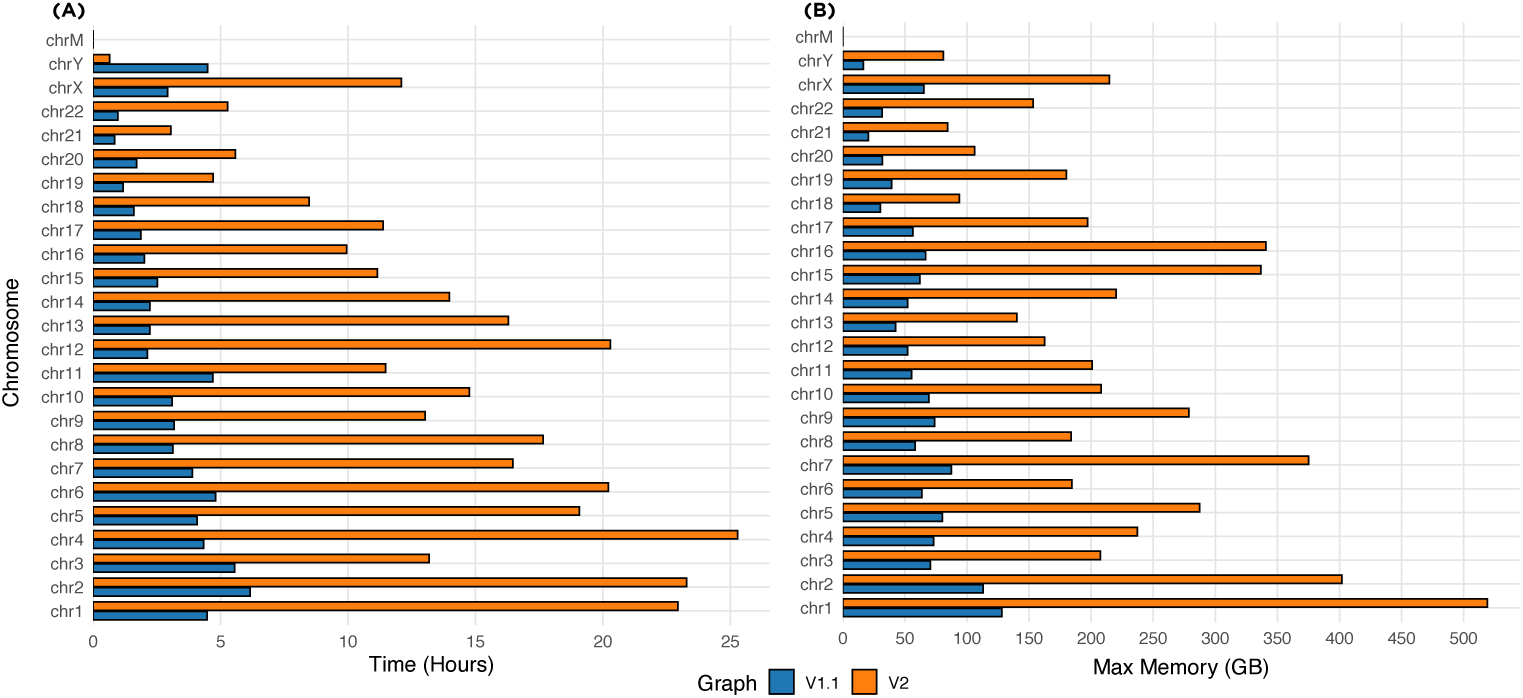
(A) is the wall-clock time, and (B) is the maximum memory usage of the tag array construction algorithm for the HPRC v1.1 and v2 graphs

After creating all the per-chromosome tag arrays, we merged them by using the whole-genome r-index that took 16 hours for HPRC v1.1 graph and 46 hours for HPRC v2 graph to create. Merging step took 7 hours on v1.1 graph and 72 hours on v2 graph. The maximum resident set size over the entire construction pipeline was 128 GiB for the v1.1 graph and 519 GiB for the v2 graph. In addition, the construction of sampled tag arrays used for coordinate translation required 104.5 GiB and 264 GiB of memory for the v1.1 and v2 graphs, respectively. The final size of the sampled tag arrays are 10-15% of the size of the full tag array index as seen in Table 1.

### 4.3 Building chromosome tag arrays across HPRC graphs

To evaluate the effectiveness of our tag array construction, we measured the proportion of BWT positions that were covered after the first two stages of tag array construction — unique *k*-mer anchoring and graph-based extension — for each chromosome sub-graph of the HPRC v1.1 and v2 graphs. These graphs include phased haplotypes from multiple individuals, with v2 representing an updated and refined version that contains 464 haplotypes, along with additional improvements in base-level accuracy and structural variant representation.

Figure 5 shows the coverage progression across all chromosomes for both versions of the HPRC graph. On the average, 95.1% of BWT positions are covered after the first two stages in v1.1, and 94.1% in v2. The coverage is particularly high in autosomes such as chr1–chr8, where repetitive content is lower and haplotype paths are more linear. In more complex or underrepresented regions such as chrX and chrY, coverage from unique *k*-mers and extensions is comparatively lower due to fewer phased samples and a higher rate of sequence redundancy.

**Fig. 5:**
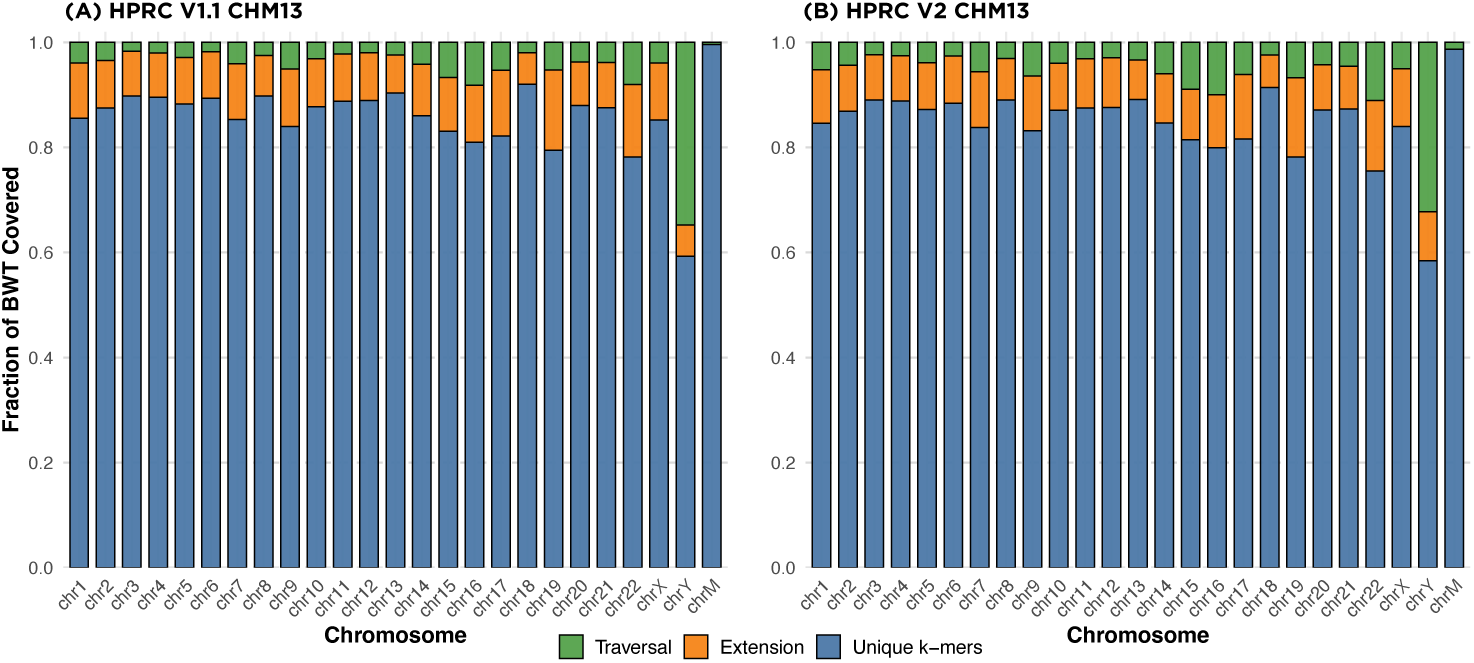
Per-chromosome tag array coverage. (A), (B) shows the coverage of each step of algorithm for the HPRC v1.1 and v2 graphs respectively

Although any remaining uncovered BWT positions are ultimately filled during the haplotype traversal stage, the results demonstrate that a large majority of the tag array can be efficiently constructed using only the initial two steps. Traversing all haplotypes in the graph is a computationally demanding process that requires substantial memory and CPU resources. The fact that nearly 95% of the tag array can be constructed using only the *k*-mer anchoring and extension steps significantly reduces the memory usage and duration of this expensive final stage. This improvement contributes to the overall scalability of the method and lowers the computational burden.

### 4.4 Scalability of building tag arrays

Figure 6(A) compares the ratio of the total number of bases, tag runs, and BWT runs between the HPRC v2 and v1.1 CHM13 graphs across chromosomes. While the total base content in v2 is roughly five times larger due to the inclusion of hundreds of additional haplotypes, the number of BWT runs increases by only about 1.2× on average, and the number of tag runs increases by about 2×. This gap demonstrates that the structural redundancy across individuals is absorbed efficiently by the index: despite the much larger sequence space, many paths remain shared, and the tag structure collapses these shared regions into longer runs. This highlights how shared sequence paths across individuals are efficiently coalesced in the tag array, minimizing duplication even as the graph becomes substantially larger and more complex.

**Fig. 6:**
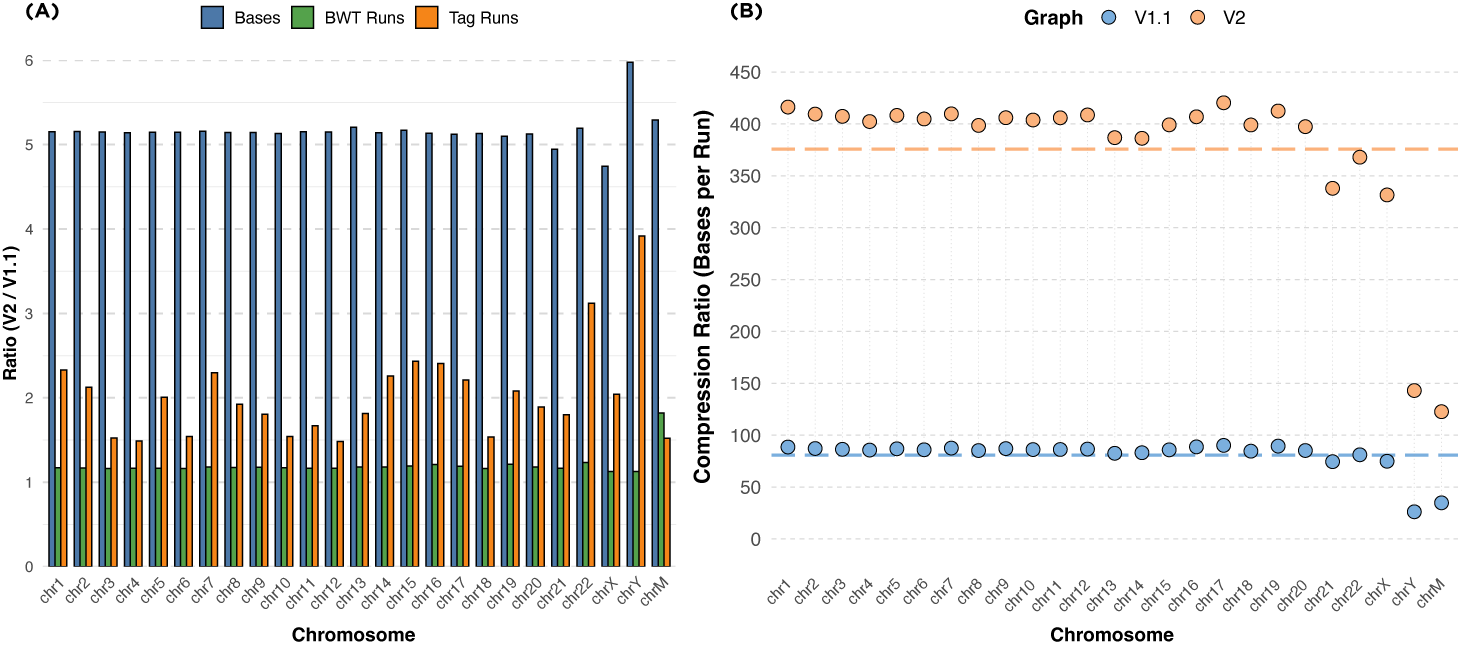
Overview of scalability and compression efficiency of tag arrays. (A) compares the ratio between number of sequence bases, BWT runs, and tag array runs between HPRC v1.1 chm13 and HPRC v2 chm13 graphs. (B) illustrates the average run-length of tags for each chromosome. The orange and blue lines shows the average run-length across all chromosomes.

Figure 6(B) further demonstrates the scalability of our method by examining the average length of tag runs in both graphs. In both v1.1 and v2, run-length encoding enables substantial compression by collapsing identical consecutive tags. However, in v2 the effect is especially pronounced, with average run lengths 375. This growth reflects both the increased redundancy from shared haplotypes and the cleaner, more contiguous assemblies present in v2. As the pangenome becomes richer and less error-prone, our method increasingly benefits from its structure. These results confirm that our tag array indexing approach scales gracefully and is highly effective for compressing and querying large, high-quality pangenome graphs.

### 4.5 Query performance

To assess the query performance of our method, we measured the time required to process 10,000 k-mers of varying lengths (from 10 to 2000), each known to appear in the BWT. For each k-mer, we first used the r-index to find the corresponding BWT interval and then queried the tag array to retrieve all unique tags within that interval. As shown in Figure 7, the total query time is highest for small k-mer sizes. In these cases, the BWT intervals are large, which leads to more unique tags to look up and makes the tag array query the dominant cost. As *k* increases, the intervals become smaller, reducing the tag array overhead, and the total query time decreases. The fastest performance is observed around *k* = 50. For larger values of *k*, the BWT intervals remain small, but the r-index query time increases due to more LF-mapping steps needed to reconstruct longer patterns. This causes the total query time to gradually rise again. These results also indicate that the assemblies in the v2 graph are cleaner. After *k* = 50, when most of the time is spent on r-index operations, the total query time on the v2 graph becomes lower than on v1.1.

**Fig. 7:**
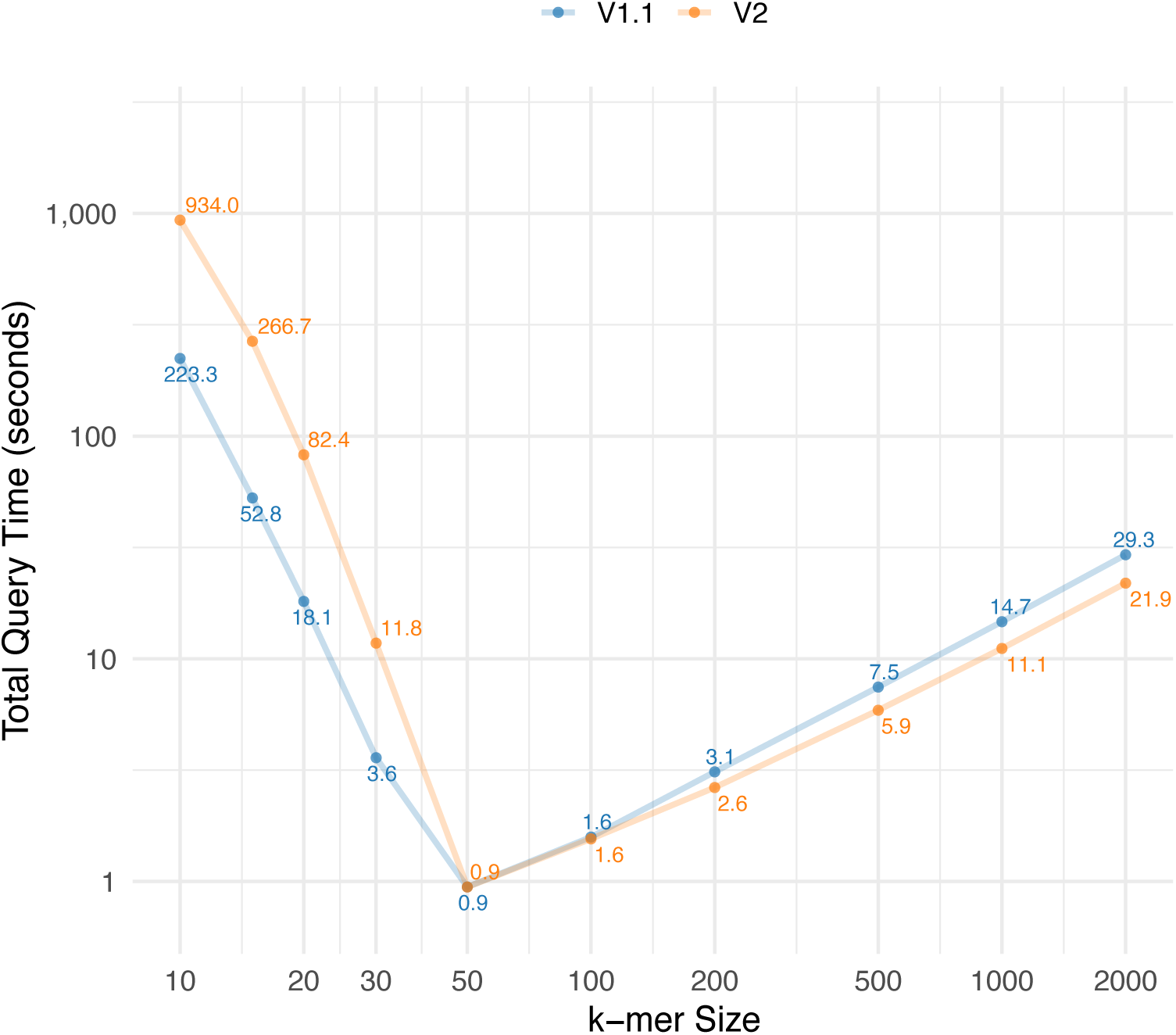
Query time for 10,000 *k*-mers with different lengths. The query time includes both the time required for r-index query and the tag array indexing query.

### 4.6 Coordinate translation performance

We evaluated the one-to-all coordinate translation queries described in Section 3.3 on the HPRC v1.1 and v2 graphs using the sampled bidirectional tag array index. For each graph, we sampled intervals of varying lengths from the haplotypes and measured the average query time as a function of the interval length. Each query was decomposed into two phases: finding the tags traversed by the input interval (find-tags) and finding all haplotypes and offsets associated with those tags (find-sequences). Figure 8 shows the total average query time across different interval lengths.

**Fig. 8:**
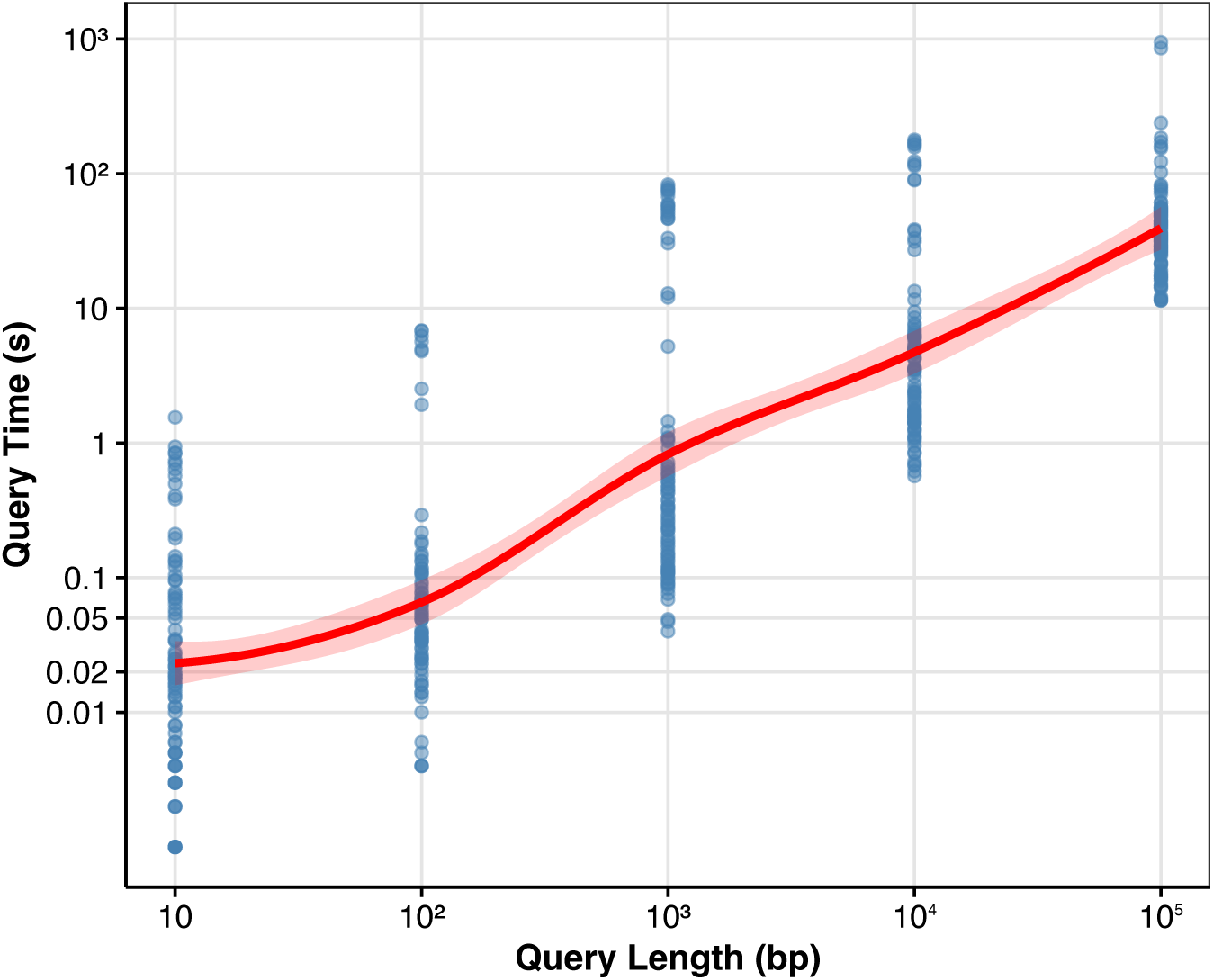
Coordinate translation query performance. Average total query time as a function of the haplotype interval length on HPRC v2 chm13 graph.

Total query time increases with interval length. Across queries with inteval length 10,000, the find-tags phase process an average of 234,152 bp/s on the v2 graph. Across all interval lengths, the find-sequences phase processes an average of 188 tags/s. The coordinate translation queries required a maximum memory usage of 104.72 GiB on the v2 graph and 71.44 GiB on the v1.1 graph.

In the find-tags phase, there is an initial overhead that scales with the length of BWT runs, while listing the tags is linear in query interval length. In the find-sequences phase, the density of tags in the query interval and the number of tag runs per tag both depend on the variation density in the region. For each tag run, there is an initial overhead that scales with the length of BWT runs, while listing the translations is linear in the length of the tag run.

### 4.7 MEM finding performance

A key application of any genomic indexing method is its ability to support fast and accurate exact-match queries. In particular, finding long MEMs [41] is critical for tasks like read mapping and variant-aware seeding. To evaluate the performance of our index in this context, we use the bidirectional extension of our method using the FMD-index. This allows extending the pattern in both directions, enabling efficient detection of MEMs. Since there are no other lossless pangenome indexes available for direct comparison, we focused our evaluation on the MEM-finding functionality and compared it to ropebwt3 [22], a highly optimized FM/FMD-index tool.

Table 2 shows the query performance of our index for MEM finding and retrieving unique tags across short and long read datasets. In most cases, the additional cost of identifying unique tags is minimal compared to total query time. This effect is especially clear for long reads and higher MEM lengths (e.g., MEM51), where BWT intervals are narrower and fewer distinct tags are associated with each match. Our MEM-finding speed is lower than that of ropebwt3, but it can be improved with further optimization. Moreover, our tag query mechanism is modular and can be integrated with any MEM-finding backend, enabling existing FM-index tools to support lossless, graph-aware mapping with minimal overhead.

**Table 2:**
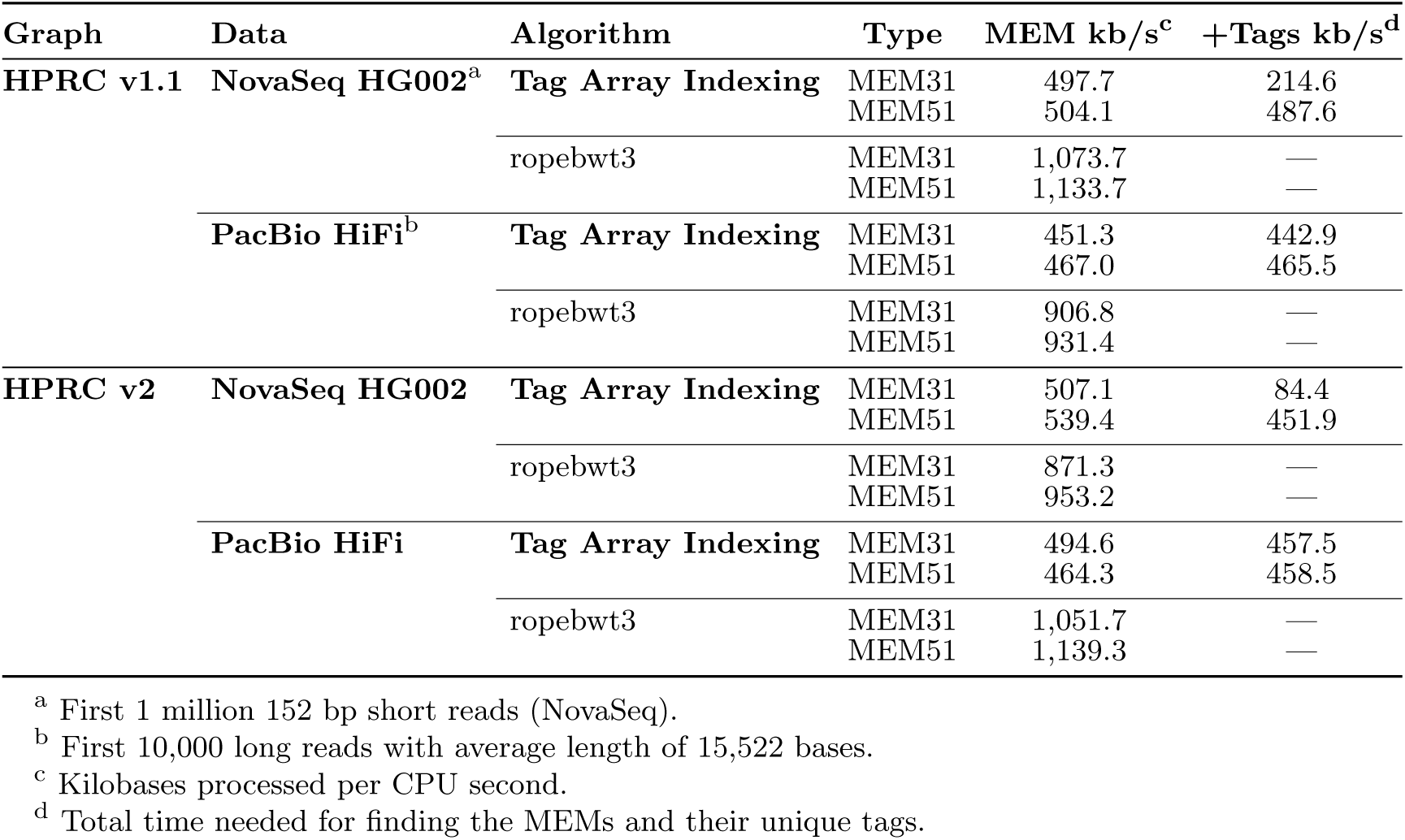
Query performance

| Graph | Data | Algorithm | Type | MEM kb/s<sup>c</sup> | +Tags kb/s<sup>d</sup> |
| --- | --- | --- | --- | --- | --- |
| HPRC v1.1 | NovaSeq HG002<sup>a</sup> | Tag Array Indexing | MEM31 | 497.7 | 214.6 |
|  |  |  | MEM51 | 504.1 | 487.6 |
|  |  | ropebwt3 | MEM31 | 1,073.7 | — |
|  |  |  | MEM51 | 1,133.7 | — |
|  | PacBio HiFi<sup>b</sup> | Tag Array Indexing | MEM31 | 451.3 | 442.9 |
|  |  |  | MEM51 | 467.0 | 465.5 |
|  |  | ropebwt3 | MEM31 | 906.8 | — |
|  |  |  | MEM51 | 931.4 | — |
| HPRC v2 | NovaSeq HG002 | Tag Array Indexing | MEM31 | 507.1 | 84.4 |
|  |  |  | MEM51 | 539.4 | 451.9 |
|  |  | ropebwt3 | MEM31 | 871.3 | — |
|  |  |  | MEM51 | 953.2 | — |
|  | PacBio HiFi | Tag Array Indexing | MEM31 | 494.6 | 457.5 |
|  |  |  | MEM51 | 464.3 | 458.5 |
|  |  | ropebwt3 | MEM31 | 1,051.7 | — |
|  |  |  | MEM51 | 1,139.3 | — |

<sup>a</sup> First 1 million 152 bp short reads (NovaSeq).
<sup>b</sup> First 10,000 long reads with average length of 15,522 bases.
<sup>c</sup> Kilobases processed per CPU second.
<sup>d</sup> Total time needed for finding the MEMs and their unique tags.

## 5 Discussion

Our tag array indexing approach represents a significant advancement in lossless pangenome indexing, overcoming key challenges in both construction and query efficiency. By combining unique *k*-mer anchoring, graph-based extension, and haplotype traversal, we achieve comprehensive coverage of the tag arrays while maintaining memory efficiency. The method’s scalability is particularly evident in the HPRC v2 graph results, where, despite a five-fold increase in base content compared to v1.1, the tag run only increased by two-fold. This demonstrates how our approach effectively coalesces shared sequences across haplotypes, allowing the index to scale sublinearly with the size of the pangenome.

The run-length encoding of tags provides substantial compression benefits, with average run lengths 375 in the v2 graph. This compression efficiency directly translates to reduced memory footprint and faster query times. Our per-chromosome construction strategy, with a divide-and-conquer-like approach, enhances scalability by enabling parallel processing and managing memory requirements for massive pangenomes. The subsequent merging step, while computationally intensive, preserves the full mapping between haplotype sequences and graph positions without loss of information.

The bidirectionaly of our method using the FMD-index opens possibilities for applications like maximal exact match (MEM) finding, which is crucial for read mapping and variant-aware seeding. While the construction time for our combined FMD-index and tag array is higher than specialized tools like ropebwt3, the additional capabilities provided by lossless graph position reporting justify this trade-off for many applications. Future work should focus on optimizing the construction pipeline, particularly the memory-intensive merging step, and exploring integration with existing read aligners to leverage the tag array’s unique capabilities for pangenome-aware alignment. Our method still requires substantial computational resources, particularly memory, with approximately 500 GiB needed to construct the bidirectional index for the v2 graph. However, given the scale of sequence data embedded in this graph, 464 haplotypes across the human genome, this memory footprint represents a reasonable trade-off. While building tag array index is memory expensive, we only need to construct the index once for each graph, and the final index requires less memory (149 GiB for the v2 graph). The construction costs are also lower than the cost of building the graphs with the Minigraph–Cactus pipeline [48].

The implementation offers configurable parameters that allow users to balance resource requirements with performance: adjusting the RLB+ tree degree and the number of runs per batch during merging creates a flexible trade-off between memory usage and CPU time, with higher tree degrees generally reducing memory requirements while increasing processing time. Several areas for improvement remain, including a faster algorithm for filling the gaps in the tag array, more efficient compression techniques for tag array storage, and optimization of the r-index structure.

Tools such as ropebwt3 build the BWT incrementally by inserting the symbols corresponding to the new sequences in the right positions in the BWT. We could similarly build the tag array incrementally by inserting new tags in the same positions. This would be a simple algorithm, not too different from the gap-filling stage of our construction algorithm. In Section 4.7, we saw that ropebwt3 uses 55–60% of the time our algorithm needs for indexing the HPRC haplotypes. By modifying ropebwt3 to build the tag array in addition to the BWT, the construction times for both algorithms would likely be similar, and the ropebwt3-based algorithm would use less memory. However, while our algorithm can reduce the wall clock time by more than a half by indexing the chromosomes in parallel, such parallelization is not possible with the incremental approach.

In addition to serving as a sequence-to-graph index, our sampled tag arrays provide a practical mechanism for coordinate translation. The r-index offers a bidirectional mapping between sequence positions and BWT positions, while the sampled tag array gives a corresponding mapping between BWT positions and graph node boundaries. Combined, these structures enable one-to-all coordinate translation: any interval on a haplotype can be mapped to all other haplotypes that traverse the same graph nodes. This allows annotations, variant intervals, and structural features to be lifted directly across haplotypes without relying on a linear reference. Moreover, because the translation operates purely on graph coordinates, the same approach can be used to relate corresponding regions across different versions of a pangenome graph.

We presented an algorithm for translating coordinates from one haplotype to all other haplotypes by computing the translation at every node in the interval. If we want to translate the coordinates to a single haplotype, a more efficient algorithm is possible. We can compute the translation at a small number of anchor nodes and traverse the haplotype paths to determine the rest. Nodes in top-level chains of the snarl decomposition [49] of the graph are particularly useful as anchors, as they are usually visited once by each haplotype.

## Appendix A Data sources

### **A.1** Graphs

- v1.1 graph: https://s3-us-west-2.amazonaws.com/human-pangenomics/pangenomes/freeze/freeze1/minigraph-cactus/hprc-v1.1-mc-chm13/hprc-v1.1-mc-chm13.gbz
- v1.1 chromosome graphs: https://s3-us-west-2.amazonaws.com/human-pangenomics/index.html?prefix=pangenomes/freeze/freeze1/minigraph-cactus/hprc-v1.1-mc-chm13/hprc-v1.1-mc-chm13.chroms/
- v2 graph: https://s3-us-west-2.amazonaws.com/human-pangenomics/pangenomes/scratch/2025_02_28_minigraph_cactus/hprc-v2.0-mc-chm13/hprc-v2.0-mc-chm13.gbz
- v2 chromosome graphs: https://s3-us-west-2.amazonaws.com/human-pangenomics/index.html?prefix=pangenomes/scratch/2025_02_28_minigraph_cactus/hprc-v2.0-mc-chm13/hprc-v2.0-mc-chm13.chroms/

### **A.2** Reads

- Novaseq HG002: https://storage.googleapis.com/brain-genomics-public/research/s equencing/fastq/novaseq/wgs pcr free/40x/HG002.novaseq.pcr-free.40x.R1.fastq.gz
- Pacbio Hifi: https://storage.googleapis.com/brain-genomics-public/research/sequencing/fastq/pacbio_hifi/HG003.1.pacbio_hifi.fastq.gz

